# Transfer learning for identification of filamentous fungi using culture and microscopy images

**DOI:** 10.64898/2026.11.10.765812

**Authors:** Susy Mathew, Jon Davis, Rebecca Davis

## Abstract

Culture morphology and microscopy images are required for identification of moulds. Automated image classification is achievable in specific domains via transfer learning of pre-trained convolutional neural networks (CNNs). The study aimed to evaluate the classification accuracy of ResNet50 CNN, for image-based classification of seven commonly isolated filamentous fungi. A total of 15,400 images were collected from seven filamentous fungi. For each species, 1,100 images each were collected for two image modalities: culture front, and microscopy, resulting in 2,200 images per class. Images from each modality for each fungus, were divided into training, validation, and test sets. The ResNet-50 model was trained and validated using four approaches: culture images (n=7,000), microscopy images (n=7,000), ensemble approach incorporating both culture and microscopy images (n=14,000), and combined culture and microscopy grid images (n=7,000). Performance was evaluated using the independent test dataset comprising images from previously unseen fungal strains, which included 700 images each, for culture, microscopy and grid images, and 1,400 images for assessment of ensemble training approach. We applied transfer learning, using both ResNet50 with and without a hierarchical approach, and evaluated their performance on the test image sets. The highest classification accuracy was achieved using the culture and grid image models, both of which achieved overall accuracy of 100%. The microscopy only and ensemble models achieved overall accuracies of 99.4% and 98.8%, respectively.

**Importance:** The decline in mycology expertise presents challenges for traditional mould identification, which relies on the interpretation of culture morphology and microscopic features. Deep learning has increasingly been applied to image classification across several areas of Microbiology and may provide an alternative approach to mould identification. In this study, we used transfer learning with a pretrained ResNet-50 CNN to classify commonly isolated filamentous fungi. We compared performance using culture images, microscopy images and approaches combining both image modalities. This work contributes to the growing evidence supporting the potential application of deep learning-based image classification for mould identification in the clinical microbiology laboratory.

## Introduction

Filamentous fungi are identified using several methods, including morphology-based identification, matrix-assisted laser desorption/ionization time-of-flight mass spectrometry (MALDI-TOF MS), and molecular methods. Traditional filamentous fungi identification in clinical mycology is primarily based on culture and microscopy morphology. However, this requires extensive mycology expertise for accurate identification. In recent years there has been a loss of mycology expertise in some Microbiology laboratories, as morphology-based identification of moulds is increasingly performed at reference laboratories due to centralisation of laboratory services. This reliance on offsite testing may create diagnostic delays. ^1^

MALDI-TOF has been increasingly utilised for identification of moulds as this method does not require extensive mycology expertise for accurate identification. However, the full protein extraction method for mould identification is cumbersome and often performed in batches, which can delay turnaround times. In addition, accurate identification depends on the availability of a comprehensive database. ^2,3^ Molecular methods like nucleic acid amplification and sequencing are currently used as complementary methods in mould identification. However, limitations like accessibility, standardisation, and relatively high cost, prevent this method from replacing traditional phenotypic testing. ^4^

Artificial intelligence (AI) is emerging as a promising tool for microbial identification. Several studies have explored the utility of computer vision in Microbiology, including the interpretation of Gram stains, identification of protozoa, and acid-fast bacilli. ^5,6,7^ Deep learning, a subset of AI, is often used in computer vision due to automated feature extraction when dealing with large data sets. More recently, deep learning has been utilized for image classification of moulds using either microscopy or culture images. ResNet-50 is convolution neural network (CNN) architecture, that has demonstrated strong performance in several studies in filamentous fungi identification. ^8,9,10^ These deep learning studies have primarily used either microscopy images or culture images for microscopy images for morphology based fungal classification. Both culture and microscopy are used by mycology experts for accurate morphology-based identification.

In this pilot study, we used transfer learning of ResNet-50 for classification of seven fungal classes using both culture and microscopy images. We also assessed the performance of ResNet-50 for culture images and microscopy images separately, as well as a hierarchical approach in training.

## Methods

### Study design and isolate selection

This single centre pilot study was conducted at the Microbiology department at NSWHP-RPA, Sydney, Australia. This was a retrospective study on stored clinical fungal isolates, obtained from clinical samples such as respiratory specimens, wound and sterile sites. Quality assurance program (QAP) isolates were also included. The clinical isolates were identified at a reference laboratory through traditional morphology-based testing. Isolates were preserved either on Sabourads dextrose agar slopes (Oxoid, Hampshire, UK) stored at 2- 8oC or in Protect microorganism preservation system (Technical service consultants, Lancashire, UK) which were stored at -80°C. Stored isolates were subcultured onto Sabourauds dextrose agar plates supplemented with antibiotics (Thermo Scientific, Thebarton, South Australia) and incubated at 25-30°C before image capture.

Seven mould species were included in the study, comprising a total of 43 strains. These included *Aspergillus fumigatus complex* (10 strains), *Aspergillus flavus complex* (6 strains), *Aspergillus niger complex* (6 strains), *Aspergillus terreus complex* (3 strains), *Scedosporium apiospermum complex* (5 strains), *Lomentospora prolificans* (5 strains), and *Penicillium* spp. (6 strains). (Table 1)

**Table 1.** Number of strains for each fungal class.

| Fungal class | Number of strains |
|---|---|
| *Aspergillus fumigatus complex* | 10 |
| *Aspergillus flavus comple* | 6 |
| *Aspergillus niger complex* | 6 |
| *Aspergillus terreus complex* | 3 |
| *Scedosporium apiospermum complex* | 5 |
| *Lomentospora prolificans* | 5 |
| *Penicillium* spp | 6 |

### Image capture

Culture images were captured using a Canon EOS R7 digital camera (Canon, Japan) fitted with a Canon EF 75-300 mm f/4-5.6 telephoto lens (Canon, Japan), which was attached using a canon EF-EOSR mount adaptor. The camera was mounted on a copy stand, and images were captured within a class II-A biosafety cabinet (MSC-advantage, Thermo Fisher Scientific, Waltham, MA, USA). Additional illumination was provided using a 5.8 W LED lamp (Daylight, London, UK). Prior to image acquisition, the plate lid was removed, and the culture plate was positioned on a white paper background. A photograph of the front of culture plate was then captured. (Figure 1) For the four *Aspergillus* and *Penicillium* species, culture images were captured between days 3-6 of incubation to encompass different stages of colony development. For *Scedosporium apiospermum complex* and *Lomentospora prolificans*, images were captured between day 5-14. To introduce variation within the image dataset, cultures were inoculated at different points on the agar plates, including one, two, three, four or more inoculation points. In addition, culture plates were rotated during image acquisition to capture colonies from multiple orientations by rotating the culture plate by 20° and capturing ten images for each orientation. A total of 100 images were captured for each plate. Wet mount preparations were made directly from colonies using the touch tape method with lactophenol cotton blue (Sigma-Aldrich, St. Louis, Missouri, USA) for microscopy. Microscopy images were captured using a Axio Lab A1 microscope (Carl Zeiss^TM^, Germany) equipped with a A-plan 40X/0.55 objective lens. A Canon EOS R7 digital camera was attached to the microscope phototube using an LM Digital SLR universal widefield adapter (LMscope, Vienna, Austria). The microscopy images were captured under standard light microscopy conditions. For both culture and microscopy, images were captured in JPEG format, using the canon EOS utility program at a resolution of 2400x1600 pixels. The images collected were deidentified without recording any patient details.

**Figure 1:**
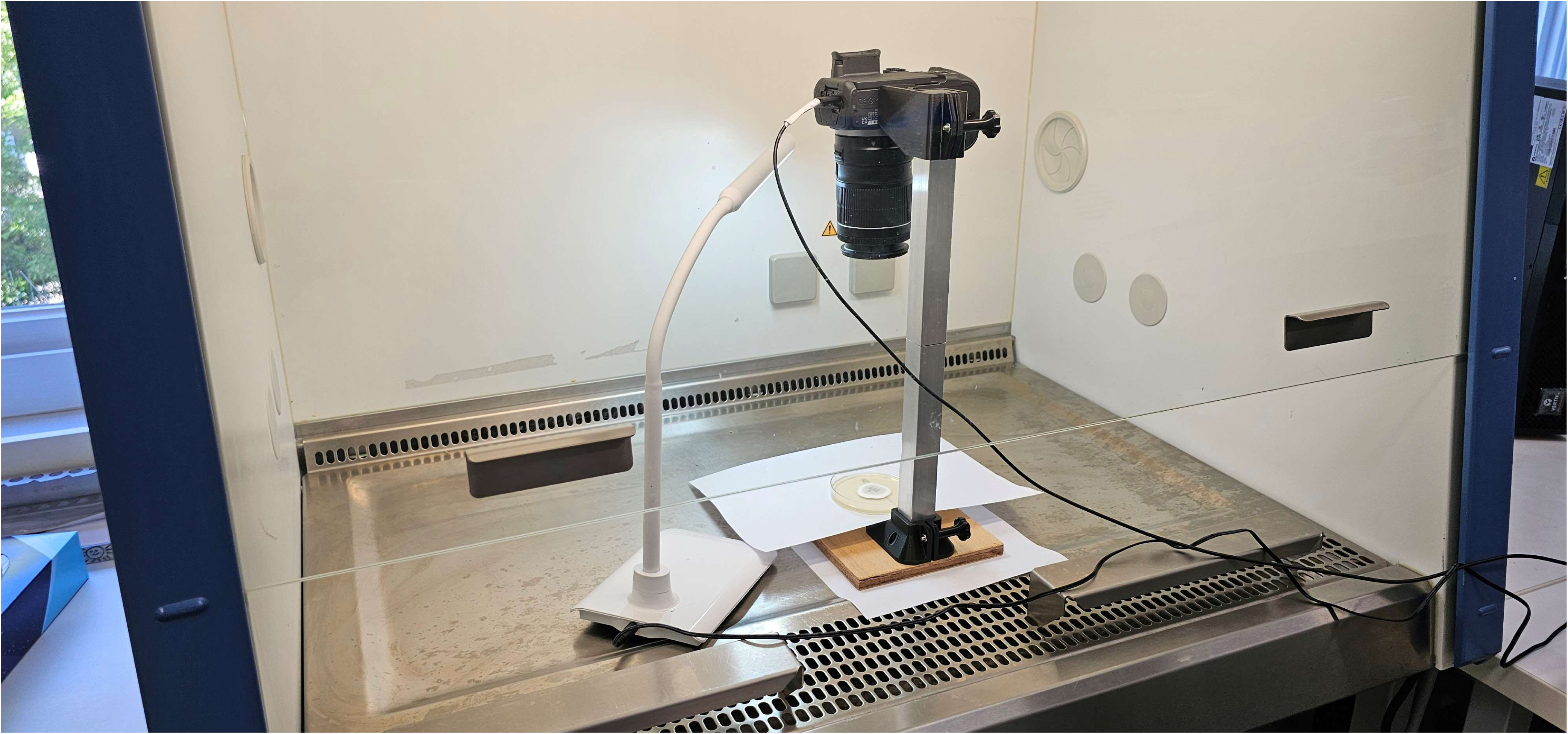
Image capture of culture plates in the biosafety cabinet.

### Image dataset

For each fungal class, 1,100 culture images and 1,100 microscopy images were collected, resulting in 2,200 images per class. A total of 15,400 images were collected for the seven fungal classes. Images were divided into training, validation and test datasets. In each class for each modality (culture or microscopy), 800 images were allocated to training dataset, 200 to the validation set and 100 images to the test dataset. Training and validation images were from the same image dataset. The test set contained images from seven previously unseen strains, with one new strain representing each fungal class. A total of 100 images were collected from each test strain, which were separate from the training and validation datasets. (Figure 2)

**Figure 2.**
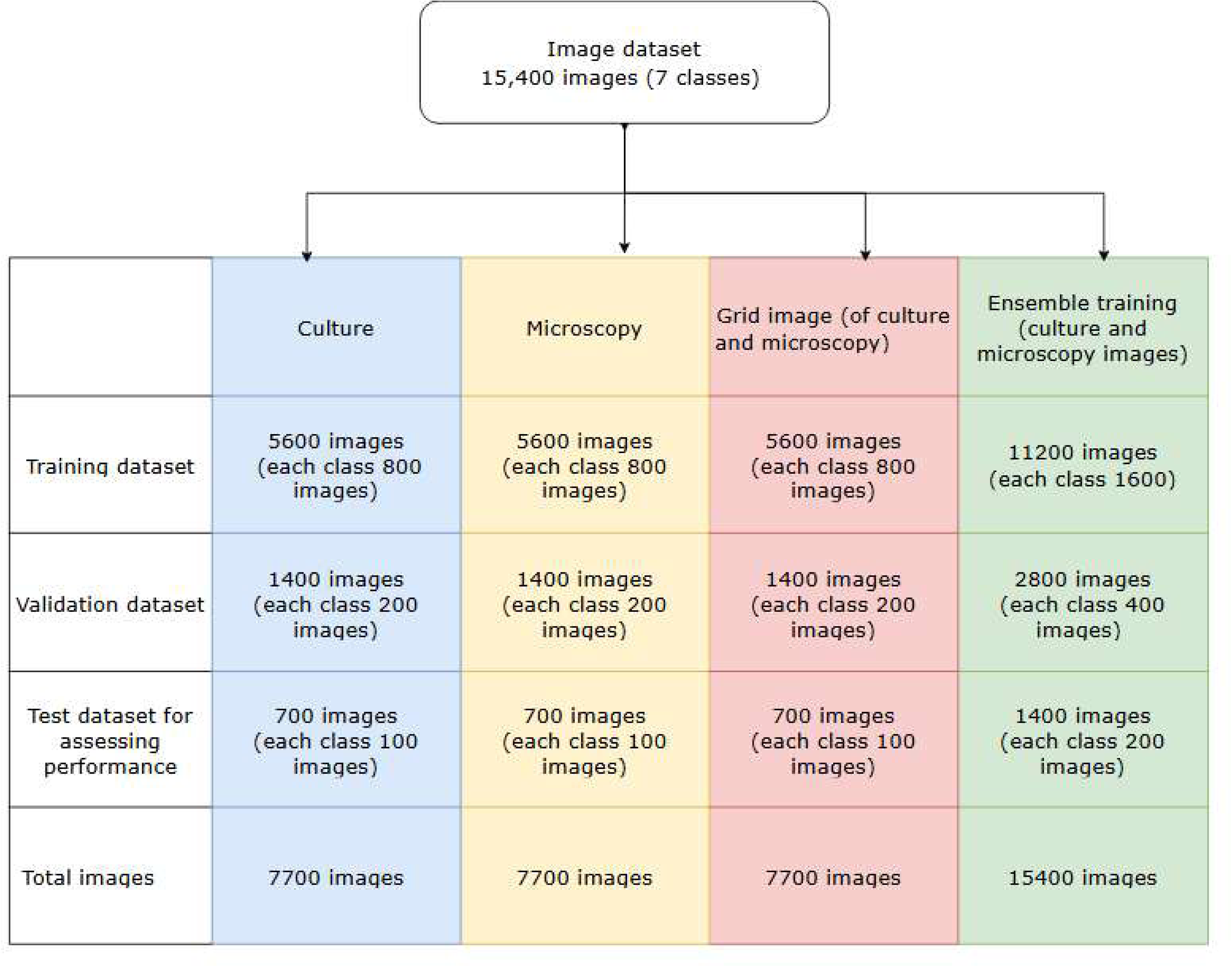
Training, validation and test dataset for culture, microscopy, grid images and ensemble training

### Deep Learning model development

Model development was performed in Google Colab using TensorFlow 2.8.0 and Python 3.12. To ensure reproducibility, a fixed random seed of 200 was used for Python, NumPy, and TensorFlow operations, and deterministic TensorFlow operations were enabled when possible. A transfer learning approach was implemented using a ResNet-50 CNN pretrained on the ImageNet dataset. Images were resized to 380x380 pixels prior to model training for macroscopy, microscopy and ensemble methods. For the grid-image approach, paired culture-front and 40X microscopy images were combined into a single composite input image with the culture image on the left and the microscopy image on the right. (Figure: 3) The resulting grid images were resized to 380X760 pixels. Data augmentation was applied during training, which included random horizontal flipping, random rotation (0.10), random zoom (0.15), and random contrast adjustment (0.15). Training was performed in two stages. In the first stage (feature extraction), the pretrained ResNet-50 backbone was frozen and used as a fixed feature extractor, while a new classification head consisting of global average pooling, dropout (0.4), and a softmax output layer was trained. The model was trained using the Adam optimizer with a learning rate of 1 x 10^-4^. In the second stage (fine tuning), the final forty layers of the ResNet-50 backbone were unfrozen and retrained to better adapt the model to mould classification. Fine tuning was performed using the Adam optimizer with a reduced learning rate of 1 x 10^-5^.

ResNet-50 was trained using macroscopy images alone, microscopy images alone, and a combined macroscopy-microscopy approach. For the combined-image approach, both ensemble training and grid-based training strategies were evaluated. In the ensemble approach, separate ResNet-50 models were trained using culture images and microscopy images using the training approach described above. Then the ensemble model combined the outputs of independently trained models to produce a final classification. For the grid-based approach, the model was trained on grid images containing both culture and microscopy images. (Figure 3)

**Figure 3:**
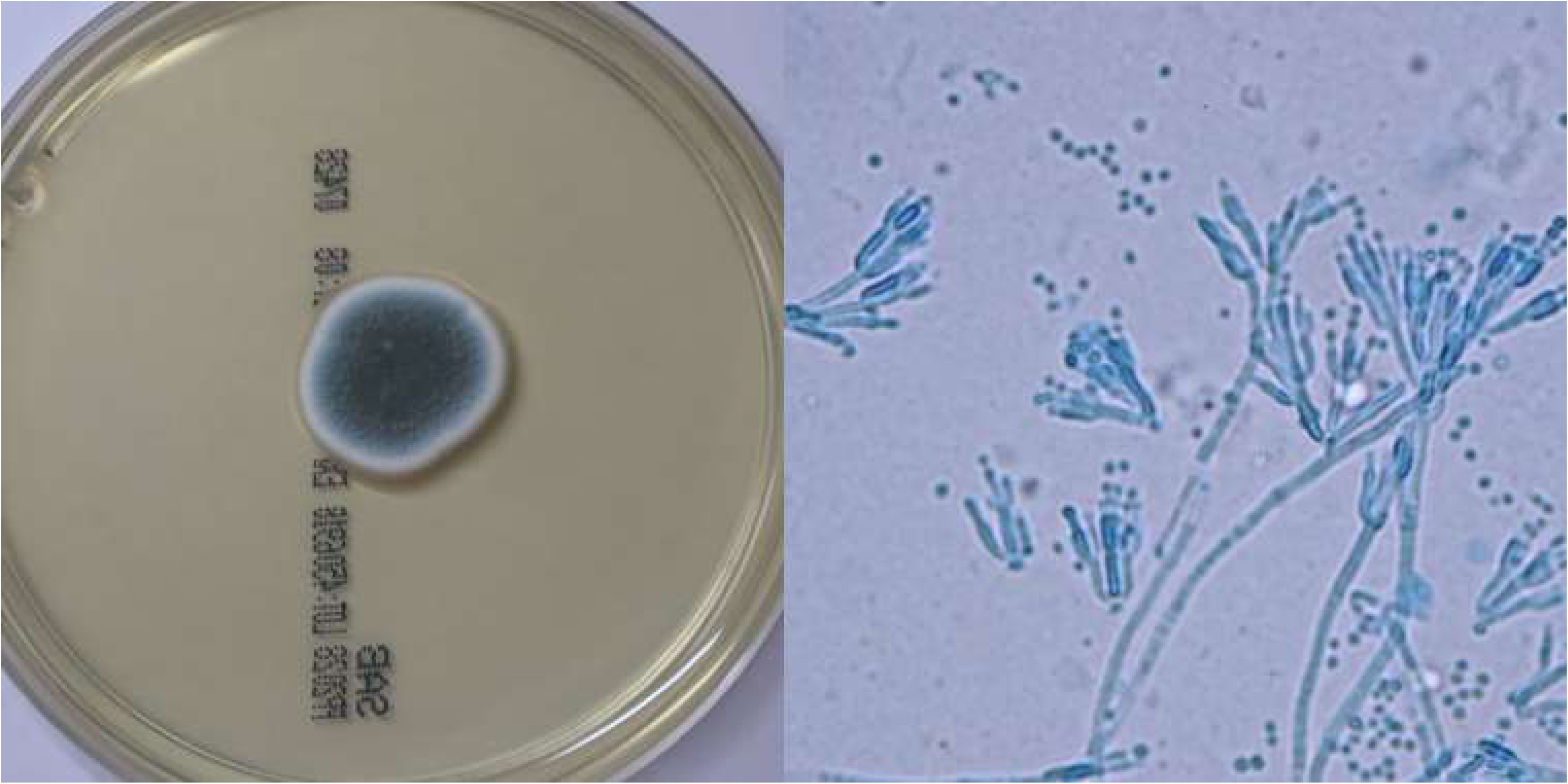
Grid image of culture plate and microscopy of *Penicillium spp*.

A hierarchical training approach was also assessed for each of the image input strategies. In this approach, an initial binary classification model was trained to distinguish *Aspergillus* from non-Aspergillu*s* isolates. Separate downstream models were trained for the *Aspergillus* group and the non-Aspergillus group to classify isolates to their respective species or species complex. The outputs of these models were subsequently combined to generate the final classification. Model training used categorical cross entropy loss and accuracy as the primary performance metric. Early stopping, model checkpointing, and adaptive learning rate reduction were used to minimise overfitting and optimise model performance. Early stopping was triggered if validation accuracy did not improve for eight consecutive epochs, while the learning rate was reduced by a factor of 0.3 after three epochs.

### Model Evaluation

Training and validation learning curves were plotted to exclude overfitting and model performance was evaluated using the independent test dataset consisting of images from previously unseen fungal strains. Performance metrics calculated on the test dataset included overall classification accuracy, plotting confusion matrices, precision, recall and F1 score. Accuracy represents the proportion of correctly classified images among all test images. Precision measures the proportion of positive predictions that were correct, while recall measures the proportion of true positive samples correctly identified by the model. The F1 score, is the harmonic mean of precision and recall, combining them to provide a balanced assessment of classification performance.

## Results

The ResNet-50 model was trained and validated on culture images(n=7,000), microscopy images (n=7,000), ensemble training(n=14,000), and combined culture and microscopy grid images (n=7,000). Performance was evaluated using the independent test dataset, which included 700 images each, for culture, microscopy and grid images, and 1,400 images for ensemble training.

The best performance was observed using culture images and grid images, both of which had an overall classification accuracy of 100%. (Figure 3 and 4) The microscopy only model and the ensemble model achieved overall accuracies of 99.42% and 98.85% respectively. (Figure 5 and 6) In the microscopy model, seven *Scedosporium apiospermum complex* isolates were misclassified as *Lomentospora prolificans,* and one *Aspergillus flavus complex* was misclassified as *Aspergillus fumigatus complex.* For the ensemble model, four *Scedosporium apiospermum complex* isolates were incorrectly classified as *Lomentospora prolificans* (96.0 % accuracy). No other misclassifications were observed for ResNet-50 without hierarchical training. Precision, recall and F1 scores were a 100% for culture and grid images, while it was 98.85% for microscopy and 99.42% for ensemble training respectively. (Table 2)

**Figure 4:**
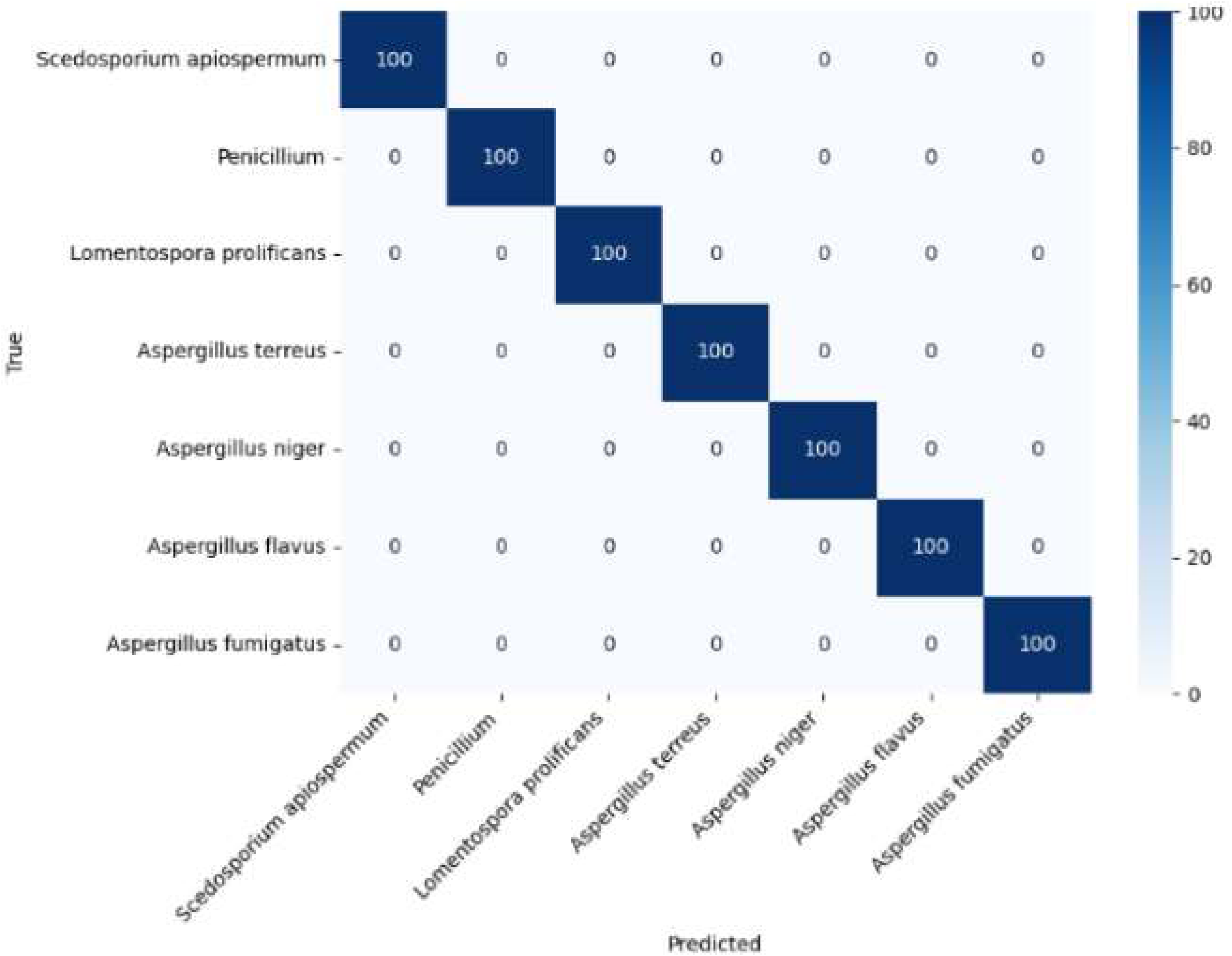
Confusion matrix for non-hierarchical training for culture images

**Figure 5:**
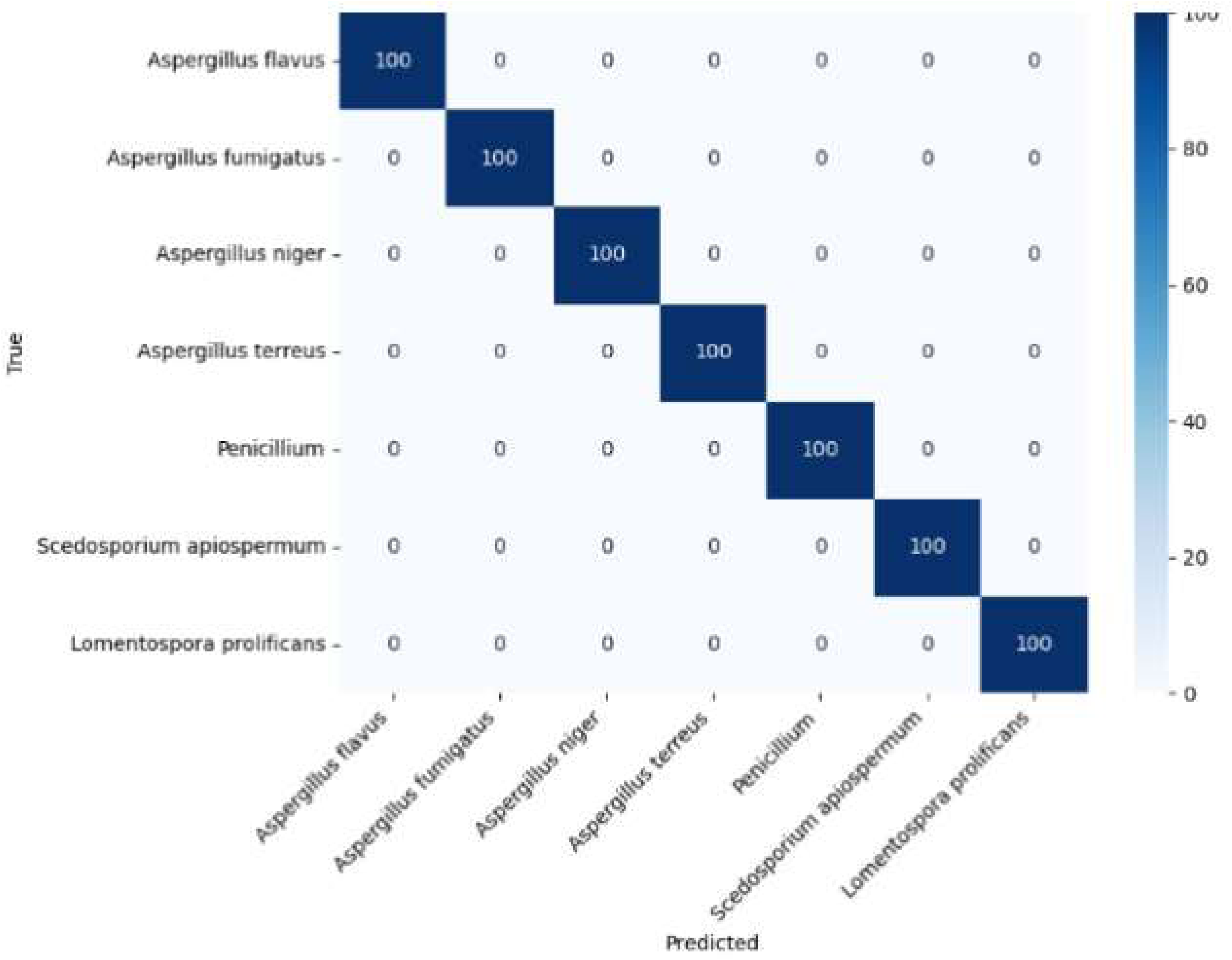
Confusion matrix for non-hierarchical training for grid images

**Figure 6:**
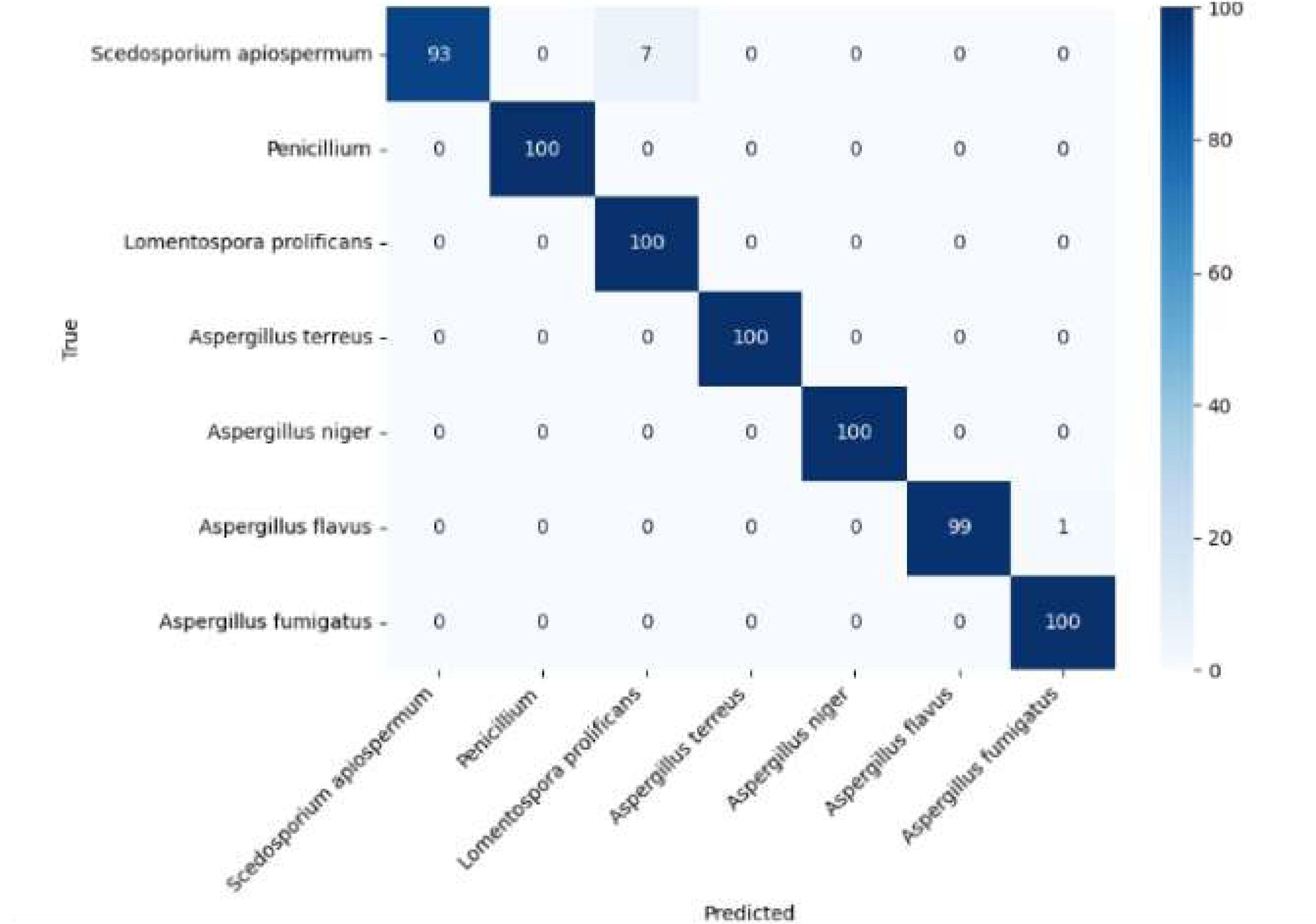
Confusion matrix for non-hierarchical training for microscopy images

**Figure 7:**
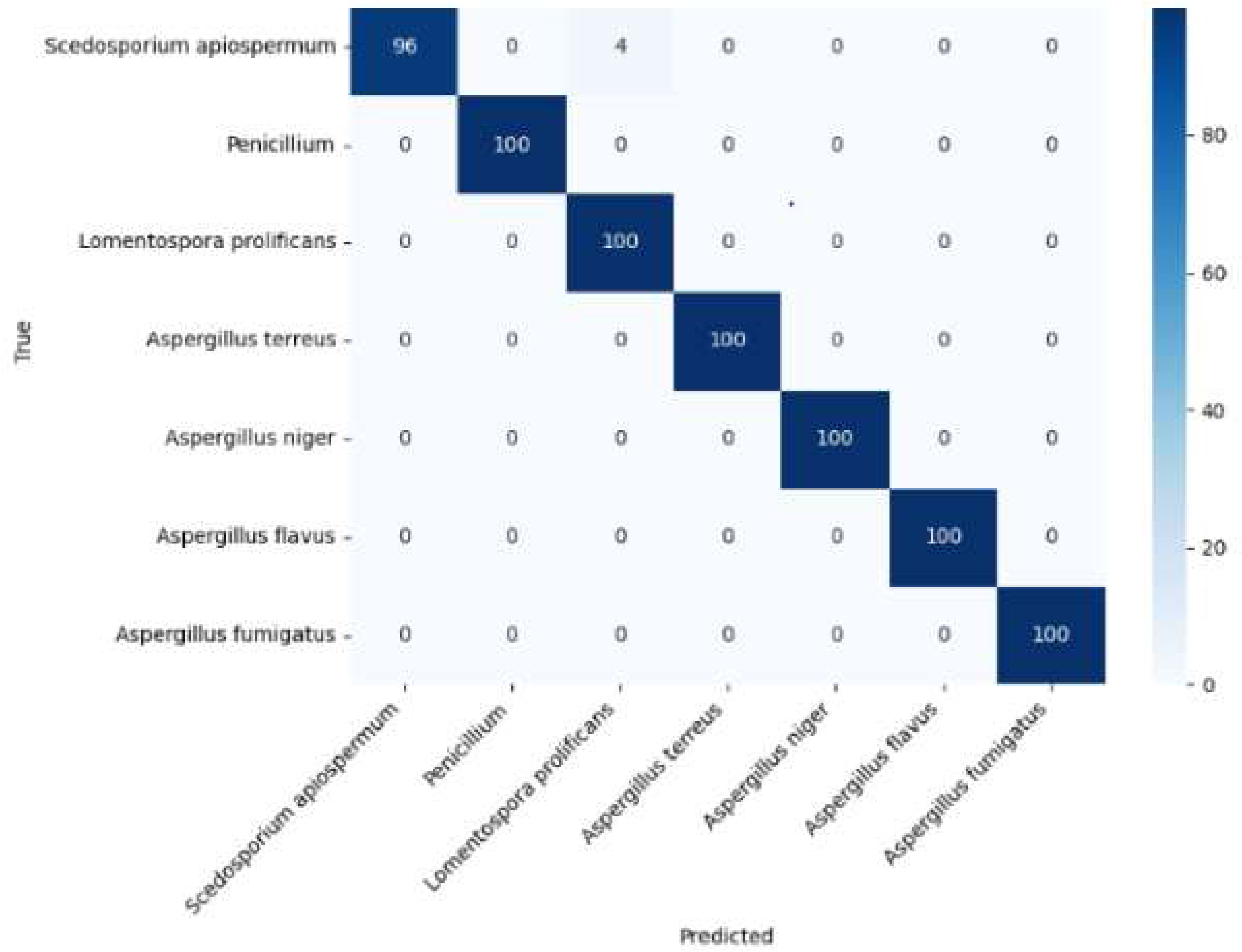
Confusion matrix for non-hierarchical ensemble training for culture and microscopy images

**Table 2.** Performance of non-hierarchical training for culture, microscopy, ensemble and grid image approach.

| Resnet50 | Culture (%) | Microscopy (%) | Ensemble approach (%) | Grid image(%) |
|---|---|---|---|---|
| Accuracy | 100 | 98.85 | 99.42 | 100 |
| Precision | 100 | 98.85 | 99.42 | 100 |
| recall | 100 | 98.85 | 99.42 | 100 |
| F1 score | 100 | 98.85 | 99.42 | 100 |
| Misclassified classes | none | Seven *Scedosporium aspiospermum complex* misclassified as *Lometospora prolificans* | Four *Scedosporium aspiospermum complex* misclassified as *Lometospora prolificans* | none |
|  |  | One *Aspergillus flavus complex misclassified* as *Aspergillus fumigatus complex* |  |  |

The hierarchical classification training approach achieved overall accuracy, precision, recall and F1 scores of 99.0% for culture, 97.85% for microscopy, 99.85% for ensemble model, and 99.40% for grid images. (Table 3)

**Table 3.** Performance of hierarchical training for culture, microscopy, ensemble and grid image approach.

| Resnet50-Hierarchical training | Culture front | Microscopy | Ensemble | Grid |
|---|---|---|---|---|
| Accuracy | 99.00 | 97.85 | 99.85 | 99.42 |
| Precision | 99.00 | 97.85 | 99.85 | 99.42 |
| Recall | 99.00 | 97.85 | 99.85 | 99.42 |
| F1 score | 99.00 | 97.85 | 99.85 | 99.42 |
| Misclassified classes | Seven *Scedosporium aspiospermum complex* misclassified as *Aspergillus terreus complex* | Two *Aspergillus flavus* misclassified ( as *Aspergillus fumigatus complex* and *Lomentospora prolificans*)<br>One *Aspergillus fumigatus complex* misclassified as *Aspergillus terreus complex*<br>Eleven *Scedosporium aspiospermum complex* misclassified as *Lometospora prolificans* | One *Aspergillus flavus* misclassified as *Aspergillus fumigatus complex* and *Lomentospora prolificans* | Four *Aspergillus flavus* misclassified as *Penicillium spp* |

## Discussion

This study utilized a pretrained CNN for the identification of seven clinically important filamentous fungi using different image input strategies. High classification accuracies were achieved across all training strategies, including accuracies for culture images and microscopy images of 100% and 98.85% respectively. Similarly high classification accuracies of 98.45% were achieved in a study conducted by Tan et al with a ResNet-50 model trained on 11,689 images from 12 *Aspergillus* species.^9^ Other studies utilising microscopy or culture images with deep learning have achieved high accuracies. ^10,11,12,13^ Our study also utilised a combined image strategy to train ResNet-50 which included an ensemble approach and grid-image approach. The grid-images achieved 100 % overall classification accuracy and the ensemble training an overall accuracy of 99.42%, with only four *Scedosporium apiospermum complex* misclassified as *Lomentospora prolificans*. To our knowledge, this is the first study which utilised both culture with microscopy images for identification of clinically relevant moulds using deep learning. A strength of this study was that model performance was evaluated using images from previously unseen strains that were not included in the training and validation datasets. The excellent performance achieved on these independent test sets, suggests that model was able to generalise beyond the isolates used during training and validation.

This study has several limitations. The number of strains included was small for some fungi, with three strains for *Aspergillus terreus complex*. Although the model demonstrated excellent performance, larger datasets with greater strain diversity and additional fungal species are needed to better assess model robustness. We hypothesize that as more fungal classes are included in training, the model may struggle to maintain high classification accuracies. Incorporating a combined culture-microscopy image input approach or a hierarchical training approach might be beneficial to improve accuracy, by providing additional discriminative features for accurate fungal identification. Another limitation was that an unclassified fungal category was not included during training. Such a category maybe significant during implementation, where the algorithm may encounter fungal species not represented in the training dataset. Hence, the performance of the current model remains uncertain when presented with fungal species not included in training. A further limitation is the absence of clinical validation as the model performance was only assessed on stored isolate. We plan to conduct a clinical validation study after training the model on more fungal classes. Despite the limitations, the results of this study are promising. The consistently high accuracies achieved across different image modalities and hierarchical training models demonstrate the potential of deep learning for mould identification. Our findings add to the growing body of literature supporting the application of AI in mould identification and suggest that deep learning may be a viable tool for early and accurate fungal identification.

## Data availability

The data sets used and analysed during the current study are available from the corresponding author upon request.

